# Integrated gut–liver-on-a-chip platform as an *in vitro* human model of non-alcoholic fatty liver disease

**DOI:** 10.1101/2020.06.10.141606

**Authors:** Jiandong Yang, Yoshikazu Hirai, Kei Iida, Shinji Ito, Marika Trumm, Shiho Terada, Risako Sakai, Toshiyuki Tsuchiya, Osamu Tabata, Ken-ichiro Kamei

**Affiliations:** Department of Micro-Engineering, Kyoto University, Kyotodaigaku-Katsura, Nishikyo-ku, Kyoto 616-8540, JAPAN; Medical Research Support Center, Graduate School of Medicine, Kyoto University, Yoshida Konoe-cho, Kyoto 606-8501, JAPAN; Institute for Integrated Cell-Material Sciences, Kyoto University, Yoshida-Ushinomiya-cho, Sakyo-ku, Kyoto 606-8501, JAPAN; Institute for Pharmacy and Molecular Biotechnology, Heidelberg University, Heidelberg, 69120, GERMANY; Faculty of Engineering/Graduate School of Engineering, Kyoto University of Advanced Science, Gotanda-cho, Yamanouchi, Ukyo-ku, Kyoto, 615-8577, JAPAN; Wuya College of Innovation, Shenyang Pharmaceutical University, Liaoning 110016, China; Department of Pharmaceutics, Shenyang Pharmaceutical University, Liaoning 110016, China

**Keywords:** non-alcoholic fatty liver disease, gut–liver axis, *in vitro* disease model, organ-on-a-chip, micro-physiological system

## Abstract

Non-alcoholic fatty liver disease (NAFLD) afflicts a large percentage of the population, but no effective treatments have been established so far because of the unsuitability of *in vitro* assays and experimental models using animals. By co-culturing human gut and liver cell lines interconnected via microfluidics for a closed circulation loop, we created a gut–liver-on-a-chip (iGLC) platform as an *in vitro* human model of the gut–liver axis (GLA) for the initiation and progression of NAFLD. Microscopic high-content analysis followed by mRNA sequencing showed that co-culturing the gut and liver cells significantly affected each cell type compared to culturing them separately. NAFLD-inducing free fatty acids (FFAs) accumulated in the gut cells and elevated gene expressions associated with retinol metabolism and glucuronidation. The FFA-treated liver cells accumulated intracellular lipid droplets and showed an increase in gene expressions associated with a cellular response to copper ions and endoplasmic reticulum stress. As an *in vitro* human GLA model, the iGLC platform may serve as an alternative to animal experiments for investigating NAFLD mechanisms.

## Introduction

Non-alcoholic fatty liver disease (NAFLD) is a common chronic liver disease that leads to hepatic steatosis, liver cirrhosis, liver cancer, and cardiovascular diseases.^1–4^ NAFLD is expected to afflict 33.5% of the United States’ population over 15 years of age by 2030.^5^ So far, liver transplantation is the only way to cure patients with severe liver disease, and finding donors that match patients is extremely difficult. While there is an urgent need to cure the different stages of fatty liver disease, the disease mechanism is largely unknown because of the complicated processes that take place at multiple layers, which is known as the multiple hit theory. For example, fat accumulation, oxidative stress, endoplasmic reticulum (ER) stress, and genetic/epigenetic modifications can take place in cells, and insulin resistance and inflammatory responses can take place in multiple organs depending on the people and environment.^6^ To find new treatments for NAFLD, a deep understanding is needed of each process, and this accumulated knowledge then needs to be combined.

In this study, we focused on the gut–liver axis (GLA), which is one of the most important components for the initiation and progression of NAFLD.^7,8^ The gut is strongly influenced by gut microbiota and diet carbohydrates and may help accelerate NAFLD.^9–11^ Inflammatory products, nutrients, and substances absorbed from food and microbiota via an intestinal barrier are carried by venous blood to the liver. In addition, products generated by hepatocytes are delivered to the small intestine. Thus, the gut and liver are close interconnected both physiologically and pathologically. GLA dysfunctions caused by NAFLD (e.g. intestinal dysbiosis, bacterial overgrowth, and alteration of mucosa permeability) are potential therapeutic targets,^12,13^ but no treatments have been made commercially available yet. This is largely because conventional preclinical animal tests do not accurately represent the problems of the multiple hit theory, lack accessibility to individual organs in living animals, and have a species difference. Therefore, establishing a simplified and robust model to study the GLA with NAFLD is crucial to obtaining deeper insights into the underlying mechanisms to discover new drugs, treatments, and diagnostic tools.

Organs-on-chips (OOCs), which are also known as micro-physiological systems (MPSs), hold great potential for *in vitro* preclinical tests^14–17^ and disease modelling.^18^ Microfluidic technology is the basis of OOCs because it allows precise control of liquid flow and three-dimensional architecture of flow channels. These properties give OOCs spatiotemporal control over cellular microenvironments to functionalise tissue cells, and a flow circulation of the cell culture medium can be generated to model multi-organ interactions with paracrine and endocrine signalling. In combination with cutting-edge cellular assays such as high-content analyses and the omics approach, OOCs provide more fundamental insights into biology in a quantitative and multi-parametric manner than animal experiments.^19^ OOCs have been applied to recapitulating the GLA *in vitro*, showing the role of crosstalk via the GLA for pathological situations (e.g. fatty liver disease^20,21^ and inflammation^22^), and for *in vitro* pharmacokinetic studies^23^. However, OOCs need to be further improved to mimic the GLA, which requires four features: a closed circulation loop, accessibility to individual chambers, dynamic flow control, and prevention of molecule absorption. The closed circulation loop is required for medium circulation to recapitulate inter-tissue interactions in the GLA. Individual accessibility is necessary to introduce tissue cells into the desired chamber and harvest them after treatment without cross-contamination from other cells. The closed circulation loop and individual accessibility seem to be contradictory features, but both need to be accomplished to investigate crosstalk between the gut and liver. Dynamic flow control is very important to obtain functional tissue cells *in vitro*, particularly for the gut.^24^. Some OOCs require the use of additional cell culture inserts (e.g. Transwell) to co-culture two or more types of cells separated by porous membrane. However, they cannot be utilised for microfabrication and thus often lack the advantages of microfluidic technology, such as control over the flow dynamics in the cell culture chamber and cellular microenvironment. Such additional inserts often interfere with microscopic observation of cells because they increase the distance between the cells and objective lens, and some porous membranes cause optical interference because of diffraction by their pores. Preventing polydimethylsiloxane (PDMS) absorption is required because PDMS is known to cause the absorption of hydrophobic molecules (e.g. metabolites, hormones, drug candidates, lipids, and fluorescent indicators), which may influence the cellular phenotypes and assay results. In particular, free fatty acids (FFAs) are a critical factor for NAFLD.

Here, we present an integrated gut–liver-on-a-chip (iGLC) platform as a simplified *in vitro* human model of the GLA, which can help with obtaining deeper insights into the underlying mechanisms of NAFLD for the development of new drugs, treatments, and diagnostic tools. The iGLA platform^25^ has micro-valves and a pump to achieve both individual accessibility for each cell culture chamber and a closed medium circulation flow to interconnect the gut and liver cells. The integrated micro-pump controls the perfusion flow to activate cultured gut cells. The iGLC platform does not require additional cell culture inserts, so it allows high-quality cell monitoring to achieve microscopic single-cell profiling. A simple surface coating with amphipathic molecules prevent the absorption of FFAs into PDMS. We co-cultured gut and liver cells with a closed circulation flow to demonstrate the viability of the iGLC platform as an *in vitro* human GLA. We also induced a NAFLD-like cellular state by administrating FFAs into the platform. Finally, we investigated unique cellular phenotypic changes and associated gene networks for the GLA in the NAFLD-like cellular state by microscopic single-cell profiling in combination with mRNA sequencing (mRNA-seq).

## Results

### Fabrication of the iGLC platform

A conceptual illustration of the iGLC platform is shown in **Fig. 1a**. The design of the microfluidic platform is a critical factor for recapitulating the GLA *in vitro*. Models based on animal experiments have difficulty with investigating the mechanism of NAFLD progression because living organs cannot be connected and disconnected to make simplified multiple-hit-theory models. To elucidate inter-tissue interaction, a single platform is needed that can both mono- and co-culture two or more types of cells within the same format. In parallel, cross-contamination needs to be considered to allow analysis of each cell type and the influence from other cell types. We designed the iGLC platform to meet these requirements (**Figs. 1b** and **1c** and **Supplementary Fig. S1**).^25^ The iGLC platform is made of PDMS with gas permeability, biocompatibility, and transparency and consists of perfusion and control layers. The perfusion layer has three sets of two cell culture chambers (2.1 mm width and 220 μm height) linked by microfluidic channels (150 μm width and 45 μm height). The control layer has a thin and flexible PDMS membrane (200 × 200 μm^2^ area and 20 μm thickness) to integrate the micro-valves and pump, which allow robust open/close control with our previously reported microfabrication technology (**Supplementary Fig. S2**).^25,26^ To investigate the interactions between two or more types of tissue cells, isolating and connecting them as desired is required. Because most OOCs do not have integrated valves and pumps, they require assembling and disassembling tubes among multiple devices, which reduces the accuracy of the control over their interactions but avoids undesired cross-contamination. The integrated micro-pump provides more accurate control than an external pump over the perfused flow within a microfluidic device. In addition, it reduces sample loss caused by extra tubing for connections. The PDMS membranes at the micro-valves and pump are actuated by the positive hydraulic pressure (150 kPa) applied by computer-controlled solenoid valves. The micro-valves allow individual cell culture chambers to be accessed without cross-contamination and precise control over sample/reagent introduction. The integrated micro-pump provides closed-loop medium circulation to recapitulate inter-tissue interactions such as the GLA and to regulate the flow rate within 0–20 nL min^-1^ by tuning of its actuating cycle.

**Fig. 1.**
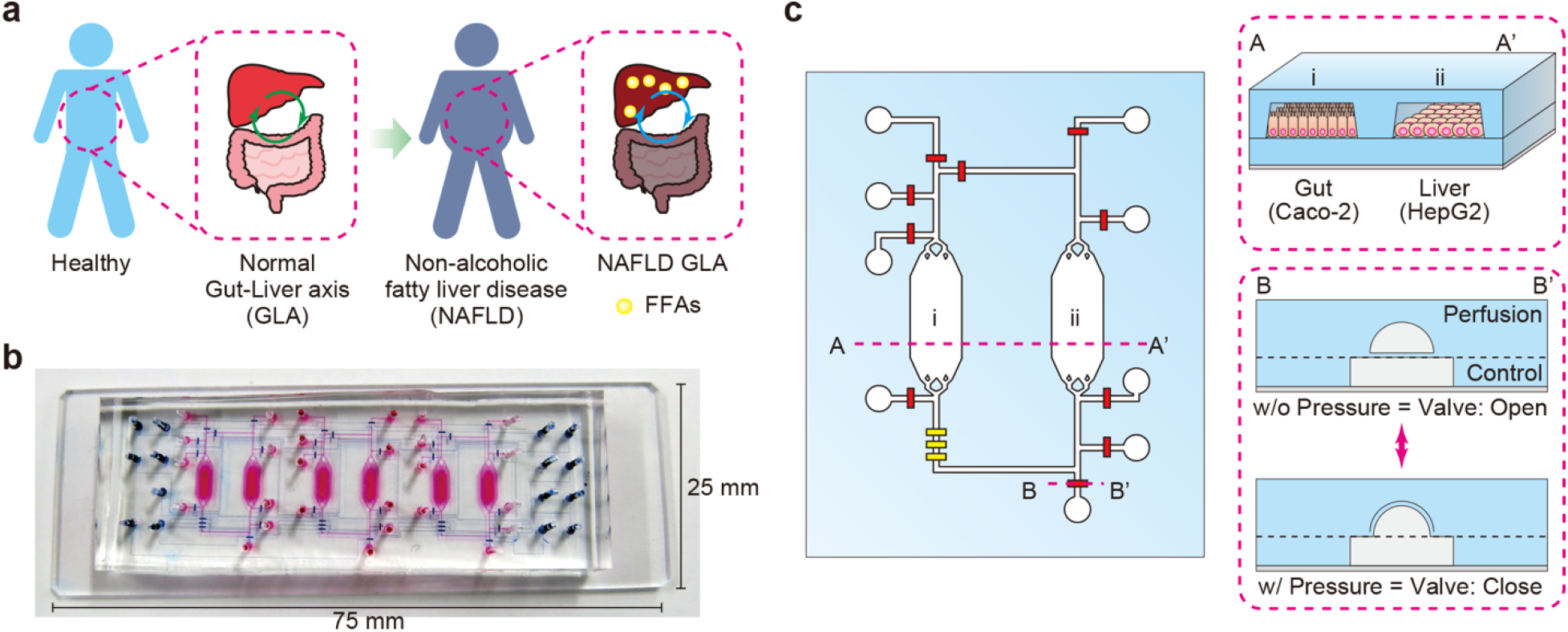
Design of the iGLC platform to recapitulate NAFLD. **a**, Schematic for NAFLD progression by FFAs via the GLA. **b**, Photograph of an iGLC platform. The perfusion and control layers are coloured in pink and blue, respectively. **c**, Illustration of the iGLC platform used for NAFLD. Two cell culture chambers are used for gut cells (Caco-2) and hepatocytes (HepG2) and linked via a microfluidic channel with a micro-pump to achieve closed medium circulation with the FFAs. With the micro-valves, each cell culture chamber is individually accessible without the risk of cross-contamination. Thus, this setup allows the inter-tissue interaction to be evaluated. A–A’ shows the cross-section of the cell culture chambers for the Caco-2 gut and HepG2 liver cells. B–B’ shows the cross-sectional view of the open and closed integrated micro valves. Because of the elastic PDMS membrane, the normally open valve is closed by applying higher pressure to the microfluidic channel in the control layer.

### Prevention of absorption of hydrophobic molecules in PDMS

Prior to the cell culture and FFA treatments, we coated the PDMS surface of the cell culture chambers with *n*-dodecyl β-D-maltoside (DDM)^27,28^ and then Matrigel to reduce the absorption of hydrophobic molecules (e.g. FFAs and AdipoRed fluorescent lipid marker)^29,30^ and increase cell adhesion and growth (**Supplementary Fig. S3**). This is a critical problem with using PDMS-based OOCs for drug discovery and *in vitro* disease modelling because small-molecule drug candidates and lipids are absorbed by the PDMS before reaching the target cells, and the concentrations differ from those with a micro-titre plate. Under such conditions, the obtained results would be unreliable. DDM is a suitable amphipathic molecule to prevent PDMS absorption because its hydrophobic part binds with the PDMS surface, and the hydrophilic part prevents the absorption of hydrophobic molecules. However, DDM is not capable of promoting cell adhesion and growth on the PDMS surface. Therefore, we tested combining DDM with a subsequent layer of Matrigel. The DDM/Matrigel coating partially prevented the absorption of the AdipoRed fluorescent lipid marker into PDMS (**Supplementary Fig. S4**). We also measured the remaining FFAs [a mixture of palmitic acid (PA) and oleic acid (OA) with the molar ratio 1:2; see Methods] in the cell culture medium after incubation with the coated PDMS (**Supplementary Fig. S5**). PA was not absorbed with both the non-coated and DDM/Matrigel-coated PDMS after 6 h of incubation at 37 °C, while it showed absorptions of 53.7% and 40.9%, respectively, after 24 h of incubation. OA showed absorptions of 22.0% and 17.4%, respectively, after 6 h of incubation, and absorptions of 67.4% and 56.7%, respectively, after 24 h of absorption. These results suggest that, although the DDM/Matrigel coating slightly mitigated the absorption of both PA and OA into PDMS after 24 h of incubation, most of the FFAs were still absorbed. Therefore, we changed the medium every 6 h during the experiments. However, further improvements need to be made to prevent PDMS absorption, or other structural materials should be used for the OOCs. This has long been a topic of discussion in this research field.^17,31^

### Modelling GLA in the chip

To recapitulate GLA *in vitro*, Caco-2 and HepG2 cells were introduced into individual cell culture chambers in the iGLC platform without cross-contamination but with a closed circulation flow (**Fig. 2a**). To confirm that the iGLC platform allows sustainable cell cultivation, the Caco-2 and HepG2 cells were cultured with a medium circulation flow of 15 nL min^-1^ for 7 days and evaluated with a Calcein AM fluorescent cell viability indicator (**Figs. 2b** and **c**). For comparison, we also tested separately cultured Caco-2 and HepG2 cells in cell culture chambers with a flow of 15 nL min^-1^ and static conditions. The flow condition did not affect the viability of HepG2 cells, which showed only some improvement with a circulation flow. The Caco-2 cells showed increased viability with a circulation flow than under the static conditions. The fluorescent intensity of Calcein AM increased by approximately 7.4-fold with a circulation flow. Caco-2 cells have previously been reported to show improved functionality and viability when cultured under continuous perfusion, which our results confirmed. This proves that the iGLC platform provides better circulation flow conditions for both Caco-2 and HepG2 cells.

We performed mRNA-seq to analyse the changes in gene expressions under the co-cultured and mono-cultured conditions and systematically investigate the crosstalk between the Caco-2 and HepG2 cells (**Supplementary Fig. S6**). A statistical test showed 159 up- and 180 downregulated genes for Caco-2 cells and 298 up- and 179-downregulated genes for HepG2 cells (**Figs. 2d** and **2e**, respectively). Based on the number of differentially expressed genes (DEGs), the HepG2 cells were more affected by co-culturing than the Caco-2 cells. The gene ontology (GO) enrichment analysis results for each gene group are shown in **Figs. 2e** and **2f.** Among the upregulated genes in the Caco-2 cells, GO terms related to the signalling pathways for cell reproduction states were found like ‘ERK1 and ERK2 cascade’ and ‘reproductive system development’. This was supported by microscopic observations (**Figs. 2b** and **2c**). Similarly, the HepG2 upregulated genes showed GO terms suggesting accelerated cell proliferation like ‘DNA replication’, ‘mitotic cell cycle phase transition’, and ‘chromosome segregation’. In addition, genes related to the fatty acid metabolic process like ACAD10, ACSM3, FABP1, and LPIN3 were included in the downregulated genes of the HepG2 cells. These results suggest that co-cultured Caco-2 cells absorb more FFAs originally contained in the cell culture medium than co-cultured HepG2 cells, which reduces the fatty acid metabolic process in the latter cells.

**Fig. 2.**
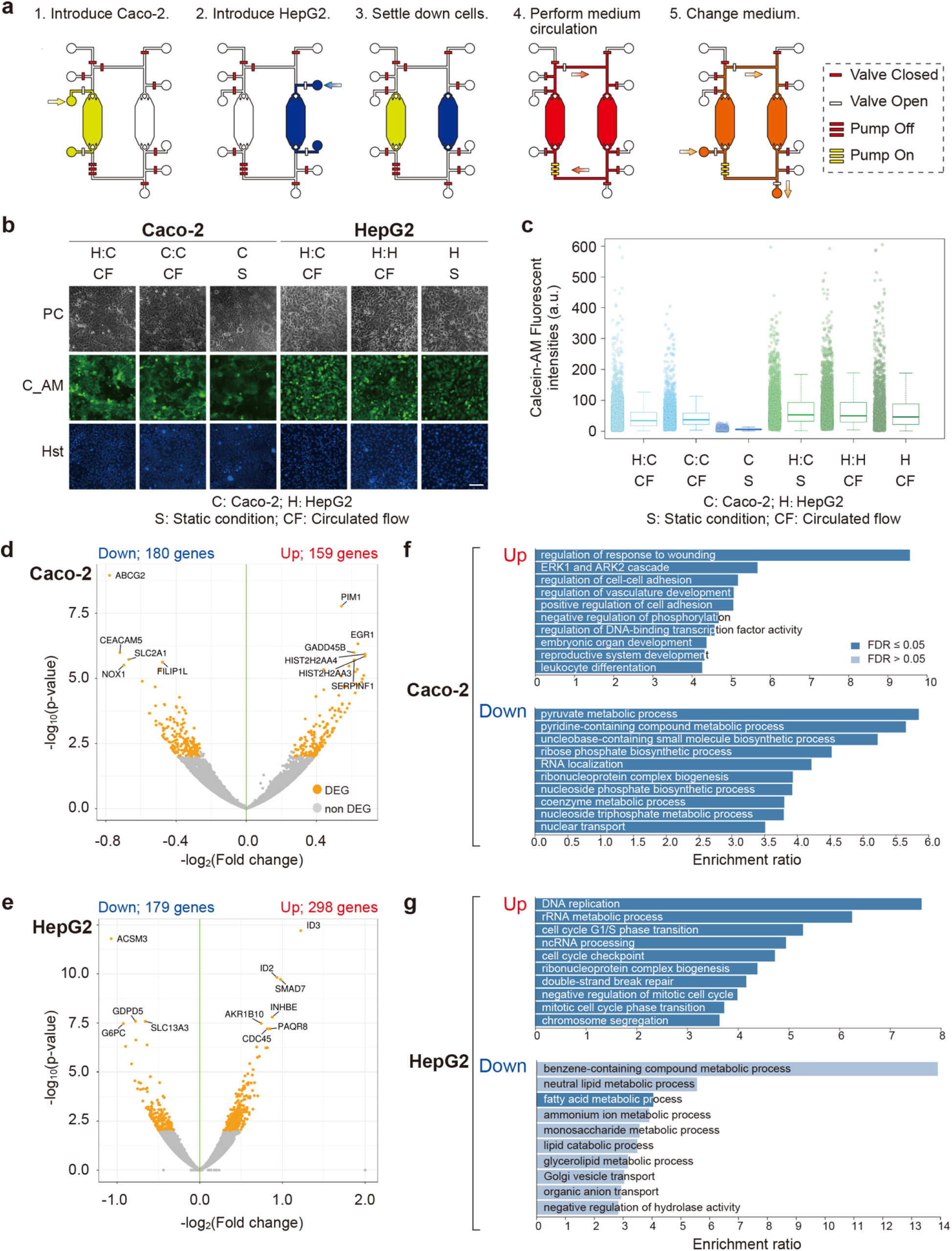
Establishment of the GLA in the iGLC platform. **a,** Experimental procedure to culture Caco-2 and HepG2 cells in an iGLC platform (see also **Supplementary Methods**). Briefly, after a microfluidic channel was washed with a fresh cell culture medium, all valves were closed to prevent air contamination into a chip. Two valves next to a cell culture chamber were opened, and cell suspensions of Caco-2 (1) and HepG2 (2) cells were separately introduced into the corresponding chambers. After 1 day of incubation at 37 °C to settle down the cells (3), the pump was actuated to circulate the medium (4). The cell culture medium was changed every 6 h (5). **b,** Phase-contrast (PC) and fluorescent micrographs of Caco-2 and HepG2 cells with closed medium circulation in the iGLC platform and stained with Calcein AM (C_AM) and Hoechst 33258 (Hst). For comparison, we also tested individually cultured Caco-2 (C:C) and HepG2 (H:H) cells under circulation flow (CF) and static conditions (S). H:C represents co-cultured Caco-2 and HepG2 cells. The scale bar represents 100 μm. **c,** Microscopic single-cell profiling of Caco-2 and HepG2 cells stained with Calcein AM. For comparison, we also tested individually cultured Caco-2 (C:C) and HepG2 (H:H) cells under circulation flow (CF) and static conditions (S). H:C represents co-cultured Caco-2 and HepG2 cells. The centrelines of the boxplots show the medians. The box limits indicate the 25th and 75th percentiles. The whiskers extend 1.5 times the interquartile range from the 25th and 75th percentiles. **d, e,** Volcano plots showing the changes in the gene expressions between the mono- and co-culture conditions for Caco-2 (**d**) and HepG2 (**e**). The top 10 genes with regard to the p-value are shown with gene symbols. **f, g,** Bar charts showing GO enrichment analysis results for up- and downregulated genes of Caco-2 (**f**) and HepG2 (**g**). The X-axis shows the enrichment ratio. Bar charts in dark blue indicate FDR ≤ 0.05 for the enrichment.

### Inducement of NAFLD on a chip

To induce NAFLD in the iGLC platform, we used FFAs represented by a mixture of PA and OA, which is typical of Western diets (**Fig. 3a**, see **Methods**).^33^ Prior to the FFA treatment, Caco-2 and HepG2 cells were cultured in the iGLC platform with serum-free DMEM for 12 h to ensure cell starvation. Then, a series of FFA concentrations from 0 to 2 mM was introduced into the iGLC platform, and the cells were incubated for 24 h with medium circulation. To evaluate the FFA accumulation in the Caco-2 and HepG2 cells, the cells were stained with AdipoRed lipid fluorescent dye to visualise the intracellular lipids (**Figs. 3b** and **3c**, **Supplementary Tables S1** and **S2**). Both Caco-2 and HepG2 cells showed intracellular FFA accumulation that was dose-dependent. The Caco-2 cells showed less FFA accumulation. Lipid droplet accumulation in hepatocytes is a hallmark of NAFLD^34^ and appeared in our model. We also carried out Annexin-V staining to evaluate the apoptotic status after FFA treatment. For comparison, the cells were also treated with 1 μM of staurosporine (STS), which induces apoptosis. Although PA accumulation has been reported to cause cytotoxicity,^35,36^ the apoptotic staining and quantitative single-cell profiling results suggested that the FFA treatment did not induce apoptosis in either the Caco-2 or HepG2 cells (**Figs. 3d** and **3e**, **Supplementary Tables S3** and **S4**).

**Fig. 3.**
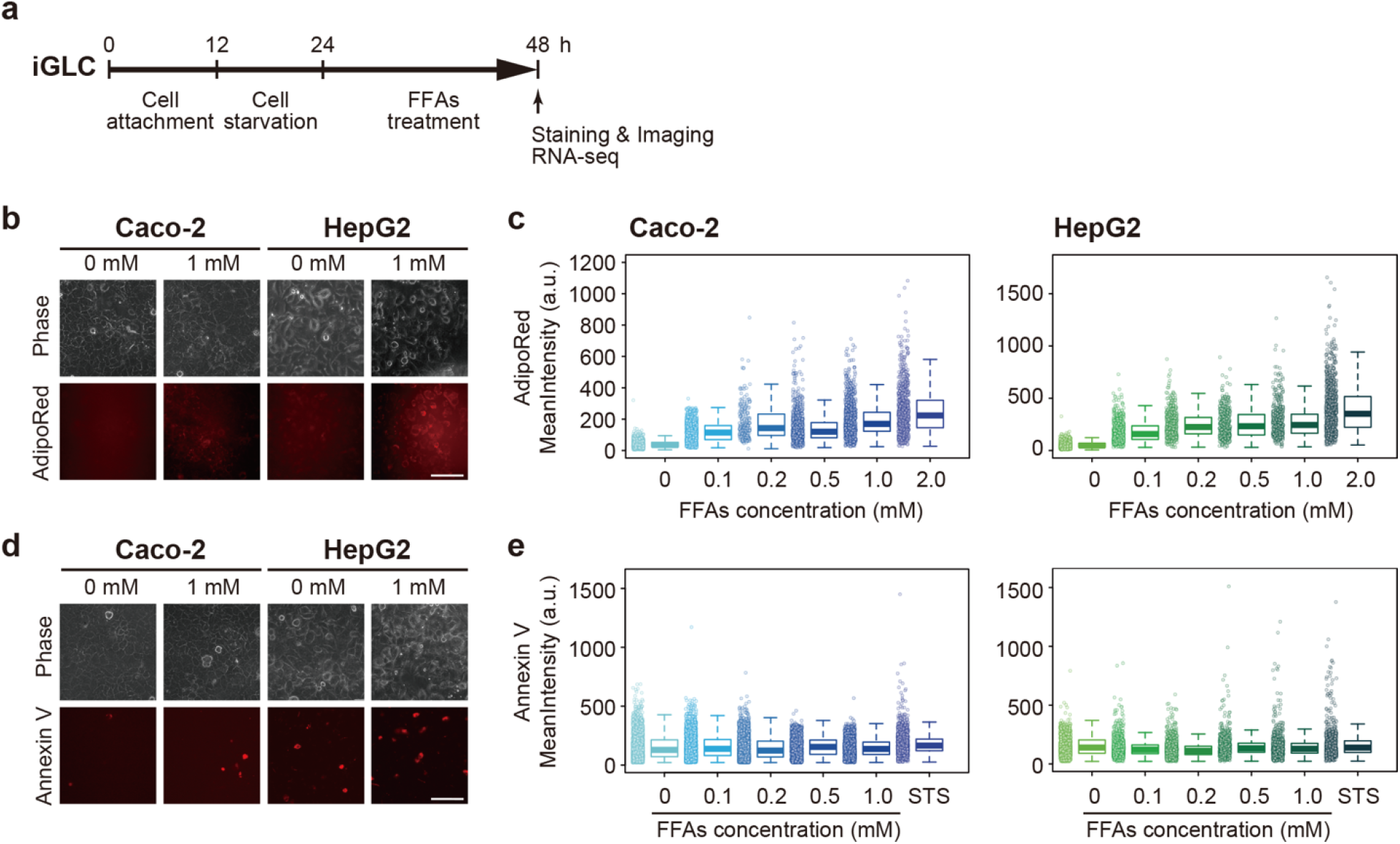
Lipids accumulated in both Caco-2 and HepG2 cells, but did not induce apoptosis. **a,** Experimental schedule to induce NAFLD in an iGLC platform. **b,** Phase contrast and fluorescent micrographs of Caco-2 and HepG2 cells treated with FFAs (0 and 1 mM) stained with AdipoRed lipid fluorescent dye. The scale bars represent 100 μm. **c,** Box plots to evaluate FFA accumulation in individual cells [Caco-2 (*left*) and HepG2 (*right*)] after FFA treatment for 24 h. The centrelines of the boxplots show the medians. The box limits indicate the 25th and 75th percentiles. The whiskers extend 1.5 times the interquartile range from the 25th and 75th percentiles. The *p*-values were estimated with the Tukey–Kramer test and are presented in **Supplementary Tables S1** and **S2** for the Caco-2 and HepG2 cells, respectively. **d,** Phase contrast and fluorescent micrographs of Caco-2 and HepG2 cells treated with FFAs (0 and 1 mM) stained with the Annexin V apoptotic cell marker. The scale bars represent 100 μm. **e,** Box plots to evaluate individual apoptotic cells [Caco-2 (*left*) and HepG2 (*right*)] after FFA treatment for 24 h. For comparison, cells were treated with 1 μM of straurosporine (STS) for 24 h. The centrelines of the boxplots show the medians. The box limits indicate the 25th and 75th percentiles. The whiskers extend 1.5 times the interquartile range from the 25th and 75th percentiles. The *p*-values were estimated with the Tukey–Kramer test and are presented in **Supplementary Tables S3** and **S4** for the Caco-2 and HepG2 cells, respectively.

To evaluate the cellular phenotypes after the FFA treatment in detail, we stained cells with Calcein AM and Hoechst 33258 fluorescent dyes. This was followed by high-content analysis (HCA) using the CellProfiler computer-guided single-cell analysis software (**Fig. 4**).^37,38^ In general, the Calcein AM and Hoechst 33258 dyes are used as cell viability and nucleus markers. However, we were unable to observe significant changes in the viabilities of Caco-2 and HepG2 cells with FFA treatment (**Figs. 4a** and **b**, **Supplementary Tables S5** and **S6**) according to the fluorescent intensity. In addition to evaluating the viability, the Calcein AM fluorescent intensity can also be applied to evaluating the cell and nucleus morphologies to obtain cellular parameters for HCA.^39,40^ Moreover, recent advances in single-cell profiling based on high-content images, such as principal component analysis (PCA) and *t*-distributed stochastic neighbour embedding (t-SNE), have allowed quantitative understanding of individual cellular status with high-dimensional cellular parameters and visualising such parameters in a two-dimensional map.^41^ We analysed microscopic images to quantify 101 cellular parameters for each cell (**Figs. 4c** and **4d**, **Supplementary Fig. S7**, and **Supplementary Table S7**) using samples of non-treated cells or cells treated with 1 mM FFA. Compared with PCA, t-SNE data visualisation allows non-treated and FFA-treated cells to be distinguished by their phenotypic changes. We found specific parameters that could be distinguished between non- and FFA-treated cells that are associated with the cellular shape (AreaShape_FormFactor, AreaShape_MeanRadius and AreaShape_MaximumRadius), Calcein AM intensity (Intensity_LowerQuartileIntensity, Intensity_MeanIntensityEdge, Intensity_MinIntensityEdge), nucleus shape (Nucleus_AreaShape_FormFactor), and Hoechst 33258 intensity (Nuclei_Intensity_LowerQuaterileIntensity, Nuclei_Intensity_MeanIntensityEdge, Nuclei_Intensity_MinIntensity and Nuclei_Intensity_MinIntensityEdge). Interestingly, the Hoechst 33258 intensity was higher in non-treated Caco-2 cells than in FFA-treated Caco-2 cells, but the HepG2 cells showed the opposite trend (**Supplementary Figs. S8** and **S9**, *p*-values in **Supplementary Tables S8** and **S9**). Thus, HCA based on simple cell and nucleus staining can be used to identify minute cellular phenotypic changes upon FFA treatment that cannot be distinguished by molecular apoptotic markers. Because the iGLC platform does not require the use of cell culture inserts, it allows such HCA-based single-cell profiling to be carried out.

**Fig. 4.**
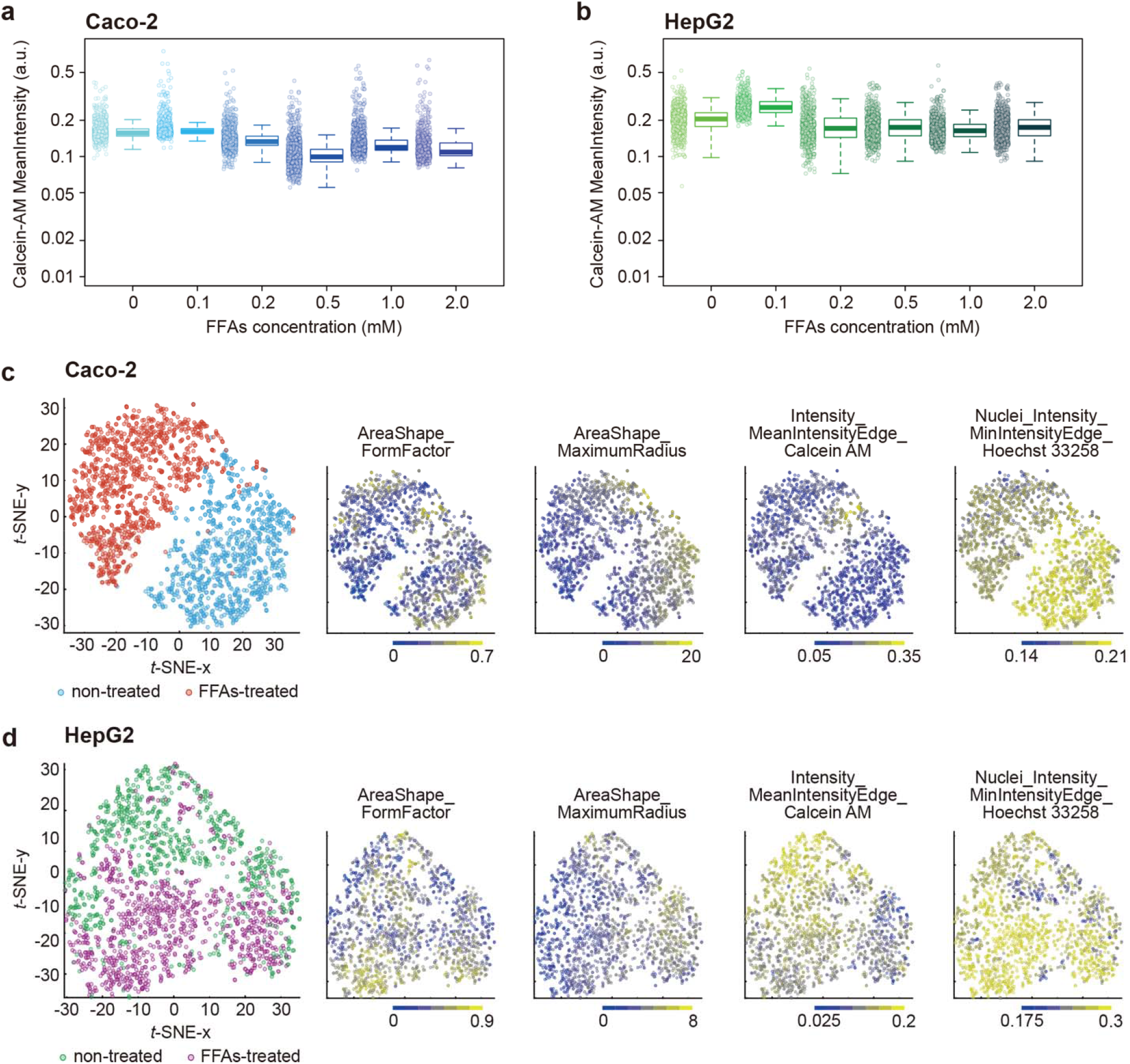
Single-cell profiling for the cell viability and t-SNE analyses. **a, b,** Box plots for the Calcein AM fluorescent mean intensities of Caco-2 **(a)** and HepG2 **(b)** cells treated with FFAs. The centrelines of the boxplots show the medians. The box limits indicate the 25th and 75th percentiles. The whiskers extend 1.5 times the interquartile range from the 25th and 75th percentiles. The *p*-values were estimated with the Tukey–Kramer test and are presented in **Supplementary Table S5** and **S6** for Caco-2 and HepG2 cells, respectively. **c, d,** Two-dimensional t-SNE plots of microscopic single-cell profiling of Caco-2 **(c)** and HepG2 **(d)** treated with 1 mM of FFAs or no treatment and stained with Calcein AM cellular and Hoechst 33258 nuclei markers. The most distinguishable cellular parameters (AreaShape_FormFactor, AreaShape_MeanRadius, and Intensity_MeanIntensityEdge of Calcein AM and Nuclei_Intensity_MinIntensityEdge of Hoechst 33258) are shown in the corresponding t-SNE plots as well as the boxplots in **Supplementary Figs. S8** and **S9**. The *p*-values are presented in **Supplementary Tables S8** and **S9**.

### mRNA-seq to identify gene expression signatures due to crosstalk for the *in vitro* GLA treated with FFAs

To elucidate the effects of the FFA treatments and the effect of crosstalk via co-culturing on the gene expressions, we performed PCA on the DEGs, which were defined by an analysis of variance (ANOVA) (see **Methods**). We found 957 and 1066 DEGs for the Caco-2 and HepG2 datasets, respectively (**Supplementary Tables S10** and **S11**). For both cell types, co-culturing effects were found with PC1 and FFA treatment (**Figs. 5a** and **5b**). The PC2 axes, which were mainly related to the FFA treatment, also showed changes in gene expression associated with co-culturing (Nos. 2 and 5 in **Figs. 5a** and **5b**, respectively) for both the Caco-2 and HepG2 cells. For the Caco-2 cells, co-culturing had the opposite effect of FFA treatment. In contrast, the HepG2 cells showed changes in the same direction along the PC2 axis with FFA treatment and co-culturing. To assess the overlapping of DEGs for each comparison, we drew Venn diagrams (**Figs. 5c** and **5d**) and heat maps (**Figs. 5e** and **5f**). We used genes showing changes in expression on the PC2 axes to focus on changes caused by the FFA treatment (see **Methods**). The Venn diagrams showed that DEGs found in mono- and co-cultured samples under FFA-minus conditions overlapped with those found in FFA-minus and -plus samples under co-culture conditions for the Caco-2 cells (**Fig. 5c**) and overlapped with those under the mono-culture conditions for the HepG2 cells (**Fig. 5d**). This suggests different types of crosstalk between co-culturing and FFAs for Caco-2 and HepG2. The heat map for DEGs showed several clusters with similar expression profiles for FFA treatments under both mono- and co-culture conditions (Clusters 2, 3, 6, and 8 in **Figs. 5e** and **5f**). We also found downregulated genes specific to the co-culture condition in Caco-2 (Cluster-1) and upregulated genes specific to the co-culture condition in HepG2 (Cluster-7).

**Fig. 5.**
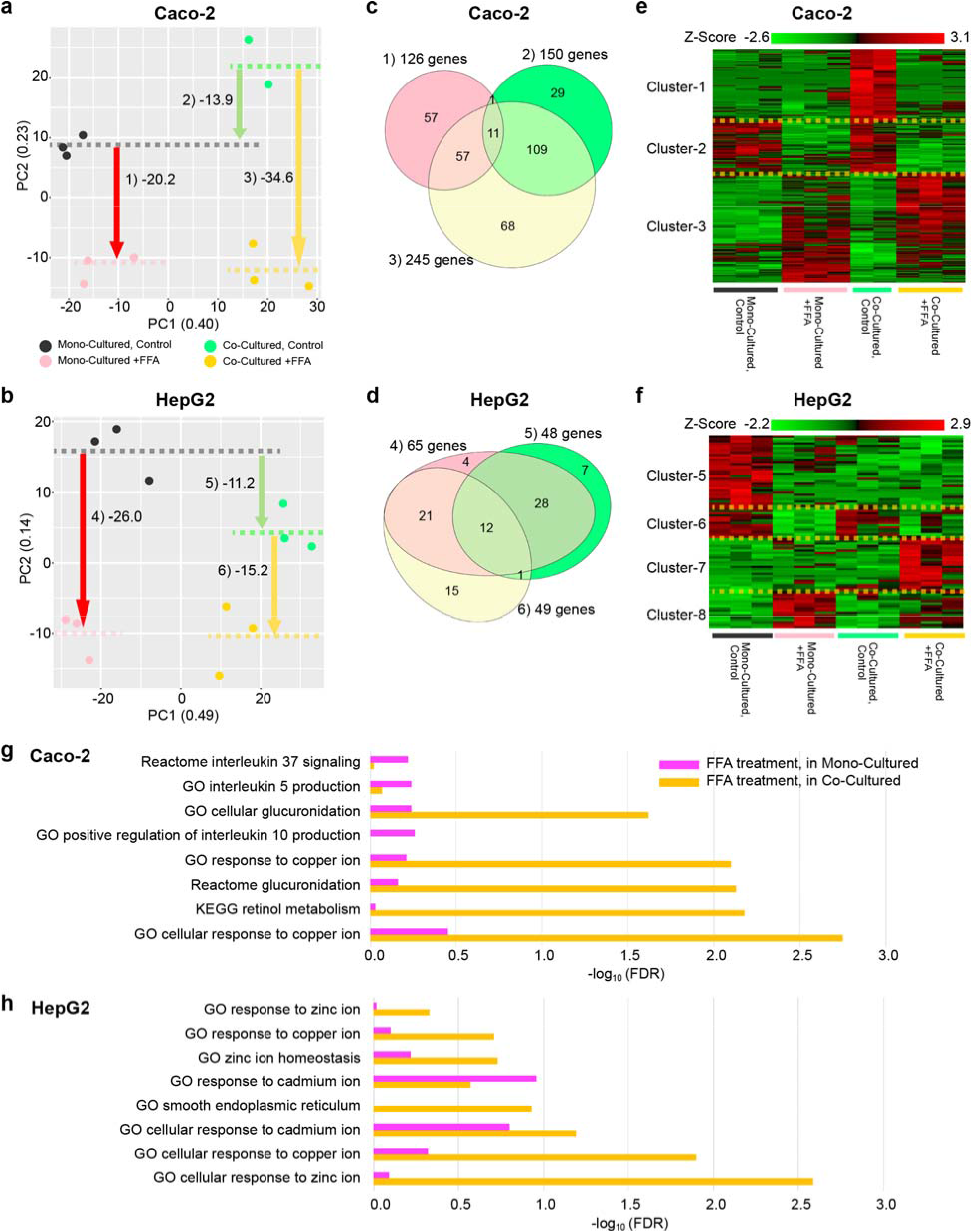
Gene expressions of the effects of FFA treatment and crosstalk with the *in vitro* human GLA model. **a, b,** PCA results for DEGs obtained from the Caco-2 (**a**) and HepG2 (**b**) experimental sets. The mean PC2 values among the replicates are shown with dotted lines. **c, d,** Venn diagram for DEGs altered according to PC2 directions (see Methods section for detail) for the Caco-2 (**c**) and HepG2 (**d**) experiments. For these diagrams, the gene name and change direction shown in **a, b** were considered. **e, f,** Heat maps for the DEGs. Z-values of the expression profiles are shown. **g, h,** Bar charts showing gene enrichment related to certain GO terms and pathways for FFA-treated Caco-2 (**g**) and HepG2 (**h**) cells under mono- and co-cultured conditions.

We then compared the effects of FFA treatment under the mono- and co-cultured conditions with gene set enrichment analysis (GSEA). This was considered with the single-cell profiling results (**Fig. 4**) to further examine the correlation between phenotypic changes in the cellular morphology with the gene regulation networks for the *in vitro* GLA in the NAFLD-like state. As noted earlier, the Hoechst 33258 intensity was slightly higher in the non-treated Caco-2 cells than in the FFA-treated Caco-2 cells (**Supplementary Figs. S8** and **S9**). The mRNA-seq results (**Fig. 5g**) did not indicate any enriched GO terms associated with transporters (i.e. ATP-binding cassette transporter ABCG2)^42^ that may efflux Hoechst 33258 outside the cells. Thus, transporters did not affect the Hoechst 33258 intensity. Interestingly, co-culturing Caco-2 cells with HepG2 cells significantly downregulated the expression of ABCG2 in Caco-2 cells (**Fig. 2d**), but this result was not influenced by FFA treatment. This suggests that FFA-treated Caco-2 cells did not efflux Hoechst 33258. As shown in **Fig. 5g**, GO terms associated with the cellular response to copper ions were enriched with an elevated gene expression. Copper is an essential trace element for human physiological processes, including enzymatic metabolism.^43^ Dysregulation of intracellular copper ions generates reactive oxygen species (ROS) and metabolic imbalance, which results in DNA damage and apoptosis.^44–46^ Our results suggest that the FFA-treated Caco-2 cells did not show the apoptotic cellular state with cellular membrane disruption but had DNA damage due to the cellular response to copper ions. The mRNA-seq results revealed that, while mono-cultured Caco-2 cells treated with FFAs showed enriched terms associated with immune systems (e.g. interleukin 5, 10, and 37 signalling), co-cultured Caco-2 cells treated with FFA showed a greatly increased number of enriched terms associated with retinol metabolism and glucuronidation (**Fig. 5g**). Retinol is an essential nutrient that needs to be obtained from the diet, and retinyl ester is delivered to organs via blood circulation by retinol binding protein 4 (RBP4). NAFLD patients show higher levels of both serum retinol and RBP4,^47^ while patients with non-alcoholic steatohepatitis (NASH) have lower levels.^48^ This indicates that the FFA-treated Caco-2 cells showed NALFD-like phenotypes rather than NASH. Glucuronidation has a major role in phase II reactions catalysed by UDP-glucuronosyltransferases (UGTs) and is involved in drug metabolism for clearing endogenous and exogenous compounds.^49,50^ UGTs are widely expressed in various organs including the liver and intestine and are also associated with drug and fat uptake and metabolism in the intestine.^51^ Although glucuronidation by UGTs has been studied with NALFD models,^52^ the role of the intestine in NAFLD is unclear. In addition, glucuronidation by UGTs shows species differences with drug metabolism.^53^ Thus, this is an important target for drug development using *in vitro* human GLA models.

In contrast, the FFA-treated HepG2 cells showed a higher fluorescent intensity of Hoechst 33258 than the non-treated cells (**Fig. 4d**). This may indicate an apoptotic cell state because Hoechst 33258 dye is often used as an apoptotic cell indicator based on an increase in the nucleus fluorescent intensity.^54^ However, the HCA results indicated that FFA-treated HepG2 underwent the apoptotic state, although this could not be elucidated by Annexin V. The mRNA-seq results revealed that HepG2 co-cultured with Caco-2 decreased the gene expression associated with fatty acid metabolic processes (**Fig. 2f**). This suggests that Caco-2 cells more effectively absorbed FFAs in the cell culture medium, so the HepG2 cells obtained fewer FFAs from the medium. Administering FFAs to the co-cultured HepG2 cells elevated cation responses such as to copper, cadmium, and zinc ions (**Fig. 5h**). Interestingly, both the FFA-treated Caco-2 and HepG2 cells showed elevated copper ions. However, the source of copper ions in the model still needs to be investigated. While NAFLD patients have lower levels of copper ions in both biopsy liver tissues and serum than healthy donors,^55,56^ NAFLD-cirrhotic patients show significantly higher levels of serum copper levels, and patients with hepatocellular carcinoma (HCC) show more elevated levels of serum copper ions.^57^ These reports indicate that copper ions have a positive relationship with NAFLD progression in cirrhotic and HCC patients. The GO term (smooth ER) was also enriched in HepG2 cells co-cultured with Caco-2 cells after FFA treatment. Smooth ER has a major role in lipid synthesis and protein processing. ER stress and the dysregulation of protein processing are strongly involved with the development and progression of NAFLD.^58–60^ Overall, the results with our platform indicate that HepG2 cells co-cultured with Caco-2 cells were activated towards NAFLD-HCC-like gene expression networks with FFA treatment.

## Discussion

We established the human iGLC platform to investigate the GLA response to FFAs for NAFLD. Although OOCs have been used to study NAFLD, the iGLC platform demonstrates several advantages based on its application of microfabrication technology. Integrating the micro-valves and pump in the platform enables both accessibility to individual cell culture chambers without undesired cross-contamination and closed medium circulation to mimic crosstalk between Caco-2 and HepG2 cells in an *in vitro* human GLA (**Fig. 2**). Because the iGLC platform does not require any additional cell culture inserts, it allows quantitative microscopic single-cell profiling. To recapitulate the NAFLD-like GLA *in vitro*, FFAs were administered in the iGLC platform. We confirmed that, although lipid accumulation occurred in both the Caco-2 and HepG2 cells, this did not induce cellular apoptosis (**Fig. 3**). We performed microscopic single-cell profiling for cell phenotypic analysis (**Fig. 4**) and mRNA-seq for global gene expression analysis (**Fig. 5**) to obtain new insights into NALFD initiation and progression.

While we were able to create the human iGLC platform, further improvements can be made for NAFLD modelling, particularly with regard to cell types. Human induced pluripotent stem cells^61^ (hiPSCs) will be helpful for the advancement of *in vitro* human NAFLD models and the iGLC platform. We were able to establish a model by using conventional and simple cell lines. However, hiPSCs generated from patients will allow their specific NALFD mechanisms to be assessed to develop precisely targeted medicine.^62,63^ There have been some cases in which patients with high-grade NAFLD had gene polymorphisms [e.g. patatin-like phospholipase-3 (PNPLA3)],^64^ and HiPSCs will allow their role in NAFLD to be studied. hiPSCs can also serve as a source to generate multi-tissue cells, which cell lines cannot. Because most cell lines like Caco-2 and HepG2 have different genomic backgrounds, they do not show proper tissue–tissue interactions. HiPSCs can also help increase the number of organs for improved *in vitro* NALFD models. Metabolic abnormality increases the risk of NALFD, type 2 diabetes mellitus, and cardiovascular diseases;^65^ to investigate their relationships, heart and pancreatic tissues can be added to the platform. Our design strategy allows the number of organs to be increased within a single device. Meanwhile, advances in gene editing technology (e.g. CRISPR-Cas gene editing)^66–68^ can be utilised not only to cure diseased hiPSCs but also to artificially generate disease-model hiPSCs.^69,70^ Recent advances in organoids from hiPSCs are also beneficial for OOC development because they allow sophisticated organ structures to be obtained *in vitro*.^7^ These cannot be performed with patients’ primary cells, which do not have sufficient long-term stability under *in vitro* culture conditions for gene editing, organoid generation, and drug testing. An iGLC platform with differentiated hiPSCs will allow pathological conditions in patients and disease models to be recapitulated for a deeper understanding of NAFLD.

Recent reports have suggested that gut microbiota significantly impact NAFLD development and should be considered for a new approach to clinical applications.^9–11^ Because the iGLC platform has individually accessible cell culture chambers, it can be used to initially culture microbiota in the gut chamber, which can then be interconnected with the liver chamber to investigate the impact of microbiota on the GLA.

In summary, the iGLC platform represents a new *in vitro* human model for recapitulating physiological and NAFLD pathological conditions with a focus on the GLA. In the near future, the iGLC platform combined with HCA and the omics approach may provide deeper insights into NAFLD development for the establishment of new drugs. The iGLC platform can contribute to the development of drugs for not only NAFLD but also a variety of disorders associated with GLA^72^ that do not have any *in vitro* experimental settings.

## Methods

### Chip fabrication

The iGLC platform was fabricated from flexible PDMS (SYLGARD 184, Dow Corning) polymer with a multilayer soft lithography replica moulding technique following the previous reports (**Supplementary Fig. S1)**.^73,74^ Briefly, the control layer consists of micro-channels supplying hydraulic pressure that were casted from a 30-μm thickness of negative photoresist mould (TMMR S2000, Tokyo Ohka Kogyo) and patterned by standard lithography. The perfusion layer consists of two cell culture chambers (225 μm in height and 2.1 mm in width) interconnected by semi-elliptical microchannels (45 μm in height and 200 μm in width). The mould fabrication process for the perfusion layer followed the multilayer lithography principle, which combines standard lithography (cell culture chambers) and greyscale lithography (microchannels). First, a negative resist layer (TMMF S2045, Tokyo Ohka Kogyo) with a thickness of 180 μm was patterned on the silicon wafer. Next, a positive resist (PMER P-LA900PM, Tokyo Ohka Kogyo) was spin-coated with a thickness of 45 μm on the wafer. Then, digital micromirror device (DMD)-based grayscale lithography (DL-1000GS/KCH, NanoSystem Solutions) was performed with numerically optimised mask data^73^ to achieve precise wafer-level mould fabrication. This allowed complete sealing of the microchannels with micro-valves and high-efficiency driving for the peristaltic micro-pumping system. After the mould fabrication, the PDMS base and curing agent were mixed well with a weight ratio 10:1. For the control layer, the PDMS mixture was spin-coated with a thickness of 50 μm on the mould to obtain a controlled PDMS thickness of 20 μm at the membrane portion. PDMS for the control and perfusion layers was cured on a hotplate at 80 °C for 4 min and in a convection oven at 80 °C for 40 min, respectively. The perfusion layer was then peeled off and precisely aligned and bonded to the control layer with the partial PDMS curing method. The assembled structure was put into the 80 °C oven for 2 h and then peeled off from the silicon wafer. Finally, the inlet and outlet wells were opened. The assembled device was permanently bonded by O_2_ plasma (FA-1, SAMCO) onto a microscopic slide glass (25 mm × 75 mm).

### Device control

The micro-valves and micro-pump were actuated by positive hydraulic pressure via linked control channels. The control channel within the chip was first filled with distilled water by a 1 mL syringe to prevent gas permeation across the PDMS. Then, metal pins and Teflon tubes (Pilot Corporation) were used to connect the inlet wells of the control channels and pneumatic system with a compressed nitrogen gas resource (regulated at 0–200 kPa). The pressure actuation and release of the valves were controlled and operated with LabVIEW (Version 11.0, National Instrument) software via solenoid valves (Microfluidic System Works Inc. and THE LEE Company) using a controller board (VC3 8 controller [ALA Scientific Instruments] and NI USB-6501 [National Instruments]). The micro-pump consisted of three adjacent micro-valves with sequential actuation to provide a periodic peristaltic motion for generating a medium recirculation flow in the chip.

### Cell culturing

HepG2 human hepatocellular carcinoma and Caco-2 human colorectal adenocarcinoma cell lines were obtained from the American Type Culture Collection. Cells were maintained with Dulbecco’s modified Eagle medium (DMEM) (Sigma-Aldrich, St. Louis, MO, USA) supplemented with 10% (v/v) foetal bovine serum (FBS, Cell Culture Bioscience), 1% (v/v) nonessential amino acids (Thermo Fisher Scientific), and 1% (v/v) penicillin/streptomycin (Thermo Fisher Scientific) in a humidified incubator at 37 °C with 5% (v/v) CO_2_. HepG2 and Caco-2 cells were passaged with trypsin/EDTA (0.04%/0.03%[v/v]) solution every 3 and 5 days, respectively, at ratios of 1/5 and 1/10, respectively.

### Cell culturing on a platform

Before cell seeding, the platform was sterilised by washing with 70% ethanol and placement under ultraviolet light in a biosafety cabinet for 30 min. Then, the cell culture chambers were coated with 0.1% (w/v) DDM (*n*-dodecyl β-D-maltoside) in PBS at 4 °C for 24 h followed by coating with Matrigel hESC-qualified matrix (Corning) diluted to 1.3% (v/v) with DMEM/F12 (Sigma-Aldrich) at 4 °C for 24 h **(Supplementary Fig. S3)**. After excess Matrigel was rinsed with DMEM, the chip was placed in an incubator at 37 °C until use.

HepG2 and Caco-2 cells were harvested from culture flasks with 1 mL of trypsin/EDTA (0.04%/0.03% [v/v]) solution and incubated at 37 °C for 5 min. After centrifuging, the cells were re-suspended in a fresh cell culture medium at 1.0 × 10^6^ cells mL^-1^. The micro-valves next to the cell culture chambers were closed to prevent cross-contamination during cell seeding. Then, 5 μL of the Caco-2 cell suspension was introduced with a pipette into the well adjacent to the cell culture chamber at 7.0 × 10^4^ cells cm^-2^. Meanwhile, 5 μL of the HepG2 cell suspension was introduced into another cell culture chamber. The platform was placed in a humidified incubator at 37 °C with 5% (v/v) CO_2_. After 1 day, the cell culture medium was changed to remove floating dead cells. The cell culture medium was then changed every 6 h under the control of our LabVIEW-based custom software.

### Free fatty acid treatment

The FFA treatment solutions were a mixture of PA (Sigma-Aldrich) and OA (Sigma-Aldrich) with a molar ratio of 1:2. To prepare the treatment solutions, PA was dissolved in dimethyl sulfoxide (DMSO; Nacalai Tesque Inc.) solution at 20 mg mL^-1^ to serve as the PA stock solution (78 mM). OA was dissolved in DMSO at 100 mg mL^-1^ to serve as the OA stock solution (354 mM). The PA and OA stock solutions were mixed (PA: OA=1: 2) in DMEM (containing 1% BSA-fatty acid free, 1% P/S, and 1 mM nonessential amino acids) to generate a series of FFA concentrations (0.1, 0.2, 0.5, 1, and 2 mM). Prior to FFA treatment, both cell culture chambers were replaced with serum-free DMEM for 12 h of cell starvation. Then, the FFA-containing medium was introduced into the cell culture chambers. The platform was incubated at 37 °C in the humidified incubator for 24 h; the medium was exchanged every 6 h.

### Cell viability and apoptosis staining

Calcein AM (Dojindo Molecular Technologies, Inc.) and Annexin V-Alexa Fluor^®^ 647 (Biolegend) dyes were used to stain viable and apoptotic cells, respectively. Hoechst 33258 (Dojindo Molecular Technologies, Inc.) was used for nucleus staining. The staining solution was prepared by mixing 10 μL of Hoechst 33258 (1 mg mL^-1^ stock concentration), 10 μL of Calcein AM (1 mg mL^-1^ stock concentration), 10 μL of Annexin V (50 μg mL^-1^ stock concentration), 500 μL of Annexin V binding buffer (Biolegend), and 500 μL of DMEM. The wash buffer comprised 500 μL of Annexin V binding buffer and 500 μL of DMEM. After treatment, cells were washed twice with fresh DMEM. Then, 10 μL of staining solution was introduced in a cell culture chamber with a pipette via the adjacent inlet. Cells were kept at 37 °C for 30 min. Then, the excess staining solution was washed away with 30 μL of washing buffer supplied three times.

### AdipoRed staining

The lipid accumulation was visualised with an AdipoRed assay (Lonza) following the manufacturer’s protocol. Briefly, 15 μL of AdipoRed assay reagent and 10 μL of Hoechst 33258 were mixed with 1 mL of DMEM to make the AdipoRed staining solution. The cell culture chambers and channels were washed with PBS twice. Then, 10 μL of AdipoRed staining solution was introduced and kept at 37 °C for 15 min. The chambers were washed with fresh DMEM solution three times.

### Image acquisition

The chips were placed on the stage of a Nikon ECLIPSE Ti inverted fluorescence microscope, which was equipped with a CFI plan fluor 10×/0.30 N.A. objective lens (Nikon, Tokyo, Japan), charge-coupled device (CCD) camera (ORCA-R2; Hamamatsu Photonics, Hamamatsu City, Japan), mercury lamp (Intensilight; Nikon), XYZ automated stage (Ti-S-ER motorised stage with encoders; Nikon), and filter cubes for the fluorescence channels (DAPI and GFP HYQ; Nikon). For image acquisition, the exposure times were set to 100 ms for (DAPI) Hoechst 33258, 5 ms for (GFP HYQ) Calcein AM, 2 s for (TRITC) Annexin V, and 100 ms for the (TRITC) AdipoRed assay.

### Ultrahigh-performance liquid chromatography-tandem mass spectrometry (UHPLC-MS/MS)

First, 3 μL of the cell culture medium was collected from a chip and then mixed with 96.95 μL of the working solution (50% isopropanol/25% acetonitrile/25% water) and 0.05 μL of the internal standard solution (PA-d4: 5 mM, OA-d9: 5 mM, in isopropanol). The mixture was then vigorously mixed for 30 s and centrifuged at 16,000× *g* for 10 min at 4 °C. Afterwards, 2 μL of the supernatant was injected and separated on a Nexera UHPLC system (Shimadzu, Kyoto Japan) by using a binary gradient with solvent A (50% Acetonitrile/18% isopropanol/32% water) and solvent B (90% isopropanol/10% acetonitrile/10 mM ammonium formate/0.1% formic acid). The gradient program was as follows: 0% B for 14 min, 100% B for 3 min, and 0% B for 3 min. An ACQUITY UPLC BEH C18 column (130 Å, 1.7 μm, 2.1 mm × 100 mm (Waters, Milford, MA)) was used at 40 °C. The UHPLC eluates were infused online to the LC-MS 8030plus (Shimadzu), which was set to negative electrospray ionisation (ESI-) mode. The PA and OA responses were observed by pseudo multiple reaction monitoring (pMRM) with transitions at m/z 255.05 > 255.35 and 281.05 > 281.45, respectively. The pMRM transitions for PA-d4 and OA-d9 were 258.95 > 259.45 and 290.10 > 290.40, respectively. The pMRM transitions were optimised, and peak areas were calculated with LabSolutions software (Shimadzu). The PA and OA responses were normalised against those of PA-d4 and OA-d9 for each sample. All measurements were obtained in triplicate, and the averaged responses were used. Standard curves were generated by measuring a blank culture medium supplemented with increasing amounts of PA and OA.

### RNA purification

RNA was purified from cells with RNeasy Mini Kit (Qiagen, Hilden, Germany). The micro-valves were initially closed to prevent cross-contamination between the HepG2 and Caco-2 cells. After the cells were washed with PBS, 10 μL of trypsin/EDTA (0.04%/0.03%[v/v]) solution was introduced into the cell culture chambers and incubated with the cells in an incubator at 37 °C with 5% (v/v) CO_2_ for 10 min. Then, cells were harvested with a 10-μL pipette and placed in 1.5-mL tubes. Cell suspensions were lysed by adding 350 μL of lysis buffer in the kit. Subsequently, 350 μL of 70% (v/v) ethanol was added in the tubes. Each solution was transferred to an RNeasy Mini spin column placed in a 2-mL collection tube. The column was centrifuged for 15 s at 8000 × *g*, and the flow-through was discarded. Then, 350 μL of buffer RW1 was added to the columns and centrifuged. Then, 80 μL of DNase digestion buffer was added to the column and incubated at 25 °C for 15 min. Next, 350 μL of buffer RW1 was added to the column tube and centrifuged again. The column was washed with 500 μL of buffer RPE twice, placed in a new 2-mL tube, and centrifuged. The column was placed in a new 1.5-mL collection tube, and 30 μL of RNase-free water was added to the column. This was followed by centrifugation for 1 min at 8000× *g* to elute RNA into the collection tube. The RNA quality was evaluated with Agilent 2100 Bioanalyser (Agilent Technologies, Inc., USA).

### RNA amplification and sequencing

RNA-seq was conducted by Takara Bio Inc. Briefly, 50 ng of total RNA from each sample was amplified and synthesised to cDNA with SMART-seq (SMART-Seq v4 Ultra Low Input RNA Kit, Takara Bio). Then, the obtained cDNA was prepared to make a library with Nextera DNA Flex Library Prep Kit (Illumina). The cDNA library was sequenced with NovaSeq 6000 (Illumina).

### mRNA-seq analysis

Initially, mRNA-seq reads were mapped to the rRNA, tRNA, or mitochondrial genome sequences with BowTie (v.2.1.0).^75^ The mapped reads were discarded and not used for the following analysis. The remaining reads were mapped to the human genome (GRCh38) with STAR Aligner (2.7.1a)^76^ using ENCODE options considering gene annotation and Ensembl ver.98.^77^ After the mapping to the genome, gene expression values (Transcripts Per Million reads; TPM) were calculated with RSEM (ver. 1.3.0).^78^ DEGs of the mono-cultured and co-cultured samples were calculated with DEseq2 (ver. 1.8.2). If a gene satisfied the following criteria, the gene was defined as DEG: p < 0.01, abs(log_2_(Fold Change)) ≥ 0.263, base mean of raw reads ≥ 31, and average TPM in either sample ≥ 1. GO analysis for DEGs was performed with the WEB-based Gene Set Analysis Toolkit (WebGestalt^80^). ‘Biological Process noRedundant’ was selected for the database, and ‘genome protein-coding’ genes were selected for the reference set. The protein-coding genes among the DEGs were used as input.

To consider the FFA treatment conditions, we employed ANOVA to select DEGs and characterise samples. A gene was treated as a DEG if p < 0.05 and abs(log_2_(Fold Change)) ≥ 0.263 for a combination of any two samples. The expression values of DEG were used for PCA to characterise the samples. To compare DEG sets with FFA-minus and -plus under the mono-cultured condition, FFA-minus and -plus under the co-cultured conditions, and mono- and co-cultured samples under the FFA-minus condition, DEGs with a PC2 loading of >0.5 or ≤0.5 were used. The results were used to assess changes in gene expression related to the FFA treatment. To compare gene expression profiles according to FFA treatment under the mono- and co-cultured conditions, we employed Gene Set Enrichment Analysis (GSEA^81^, ver.4.0.4) with DB files (msigdb.v7.1.symbols.gmt). Among the results, we selected the terms of ‘Copper ion’, ‘Retinol’, ‘Glucuronidation’, and ‘Interleukin’ for Caco-2 analysis and the terms of ‘Copper ion’, ‘Zinc ion’, ‘Cadmium ion’, and ‘Smooth endoplasmic reticulum’ for the HepG2 analysis. The fold change values for the protein coding genes were used as inputs. The bar plots show FDR q-values for the top four terms under either the mono- and co-cultured condition. In the analysis, the volcano plots and PCA plots were drawn with the ggplot2 package in R.

### Single-cell profiling based on microscopic images

Following the microscopic image acquisition, the CellProfiler software (Broad Institute of Harvard and MIT, Version 3.1.9) was used to identify cells with Otsu’s method. The fluorescence signals of individual cells were quantified automatically. Further analysis of single-cell morphological descriptors was performed with PCA and t-SNE in the open-source Orange 3 software (Version 3.23.1; Bioinformatics Laboratory, Faculty of Computer and Information Science, University of Ljubljana, Slovenia).^82^

### Statistical analysis

The Tukey–Kramer test and asymptotic Wilcoxon signed rank test were carried out with *R* software (ver. 3.5.2; https://www.r-project.org/).

### Data availability

The mRNA-seq data have been deposited in the NCBI Gene Expression Omnibus (GEO) under accession number GSE152091.

## Acknowledgements

We thank Ms. Miyako Fujita for maintaining the cells. Funding was generously provided by the Japan Society for the Promotion of Science (JSPS) (16K14660, 17H02083, 18KK0306, and 19H02572), the Terumo Life Science Foundation, the Ebara Hatakeyama Memorial Foundation, Japan Agency for Medical Research and Development (AMED) (17937667) and LiaoNing Revitalization Talents Program (XLYC1902061). MT was supported by the Nakatani Foundation for the Advancement of Measuring Technologies in Biomedical Engineering. Part of this work was supported by the Nanotechnology Platform Project within MEXT, Japan, through the Kyoto University Nano Technology Hub. WPI-iCeMS is supported by the World Premier International Research Centre Initiative (WPI), MEXT, Japan.

## Author contributions

J.Y., Y.H., T.T., O.T., and K.K. conceptualised the work. J.Y. and Y.H. fabricated the device. J.Y., R.S., and S.T. performed the cell culture experiments and analyses. S.I. carried out the UHPLC-MS/MS experiments. J.Y., R.S., S.T., and K.K. performed the image analysis. J.Y., S.T., and M.T. performed the RNA experiments. I.K. and K.K. performed the RNA-seq analyses. All authors contributed to data analysis, discussion, and interpretation. J.Y., Y.H., and K.K. wrote and revised the manuscript with input from all authors.

## Supplementary Information

**Supplementary Fig. S1.**
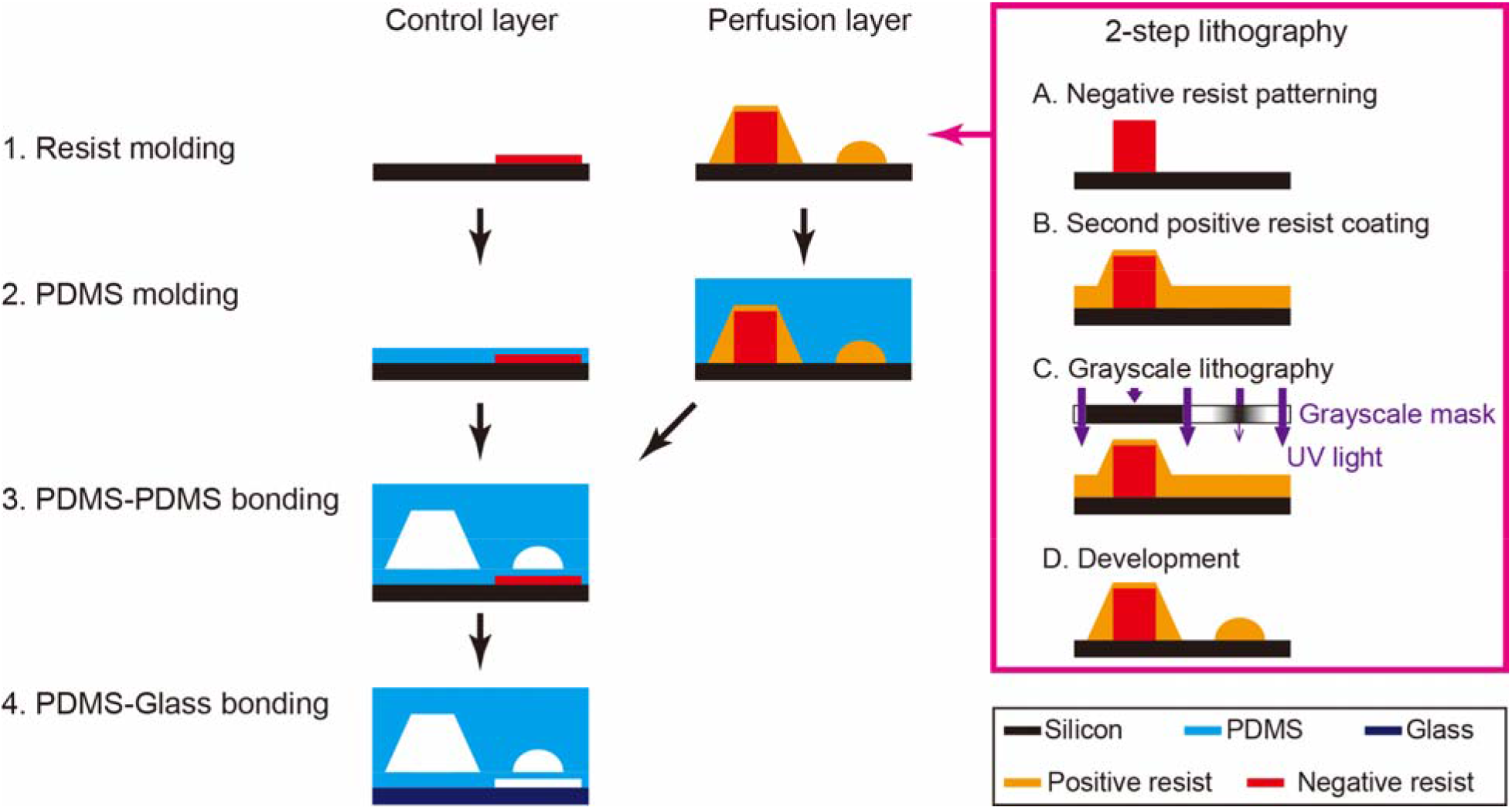
Fabrication procedure for the iGLC platform.

**Supplementary Fig. S2.**
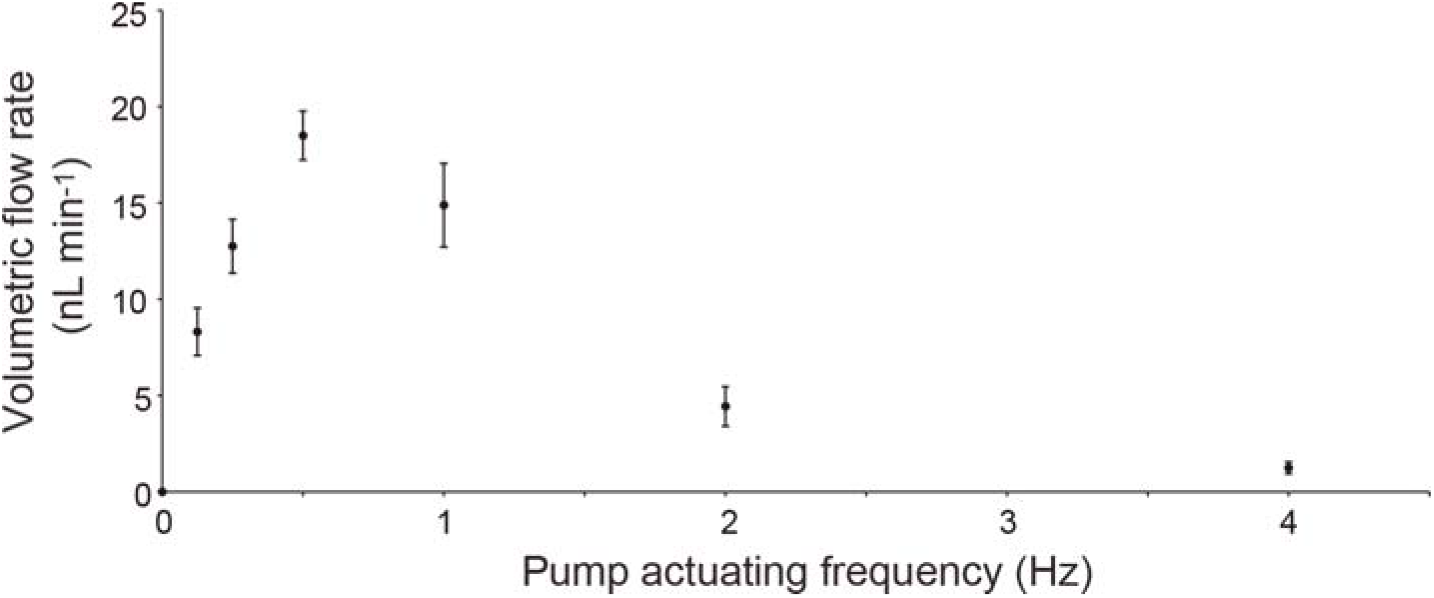
Flow rates of the cell culture medium in the iGLC platform operated by the integrated micro-pump. The flow rates were regulated by the actuation frequency for the sequential opening and closing of a set of three micro-valves acting as a micropump. To evaluate the flow rates, micro-beads (4.5 μm in diameter; Polysciences, Inc.) were used to visualise the medium flow and measure the flow distance. Prior to measurements, the microfluidic channel was coated with 1% (w/v) bovine serum albumin (BSA, Sigma-Aldrich) in PBS for 2 h at 25 °C to prevent non-specific adhesion on the channel. Micro-beads were suspended in 1% (w/v) BSA solution at 1.0 × 10^6^ beads mL^-1^. Then, 5 μL of bead-containing solution was introduced into the chip. After a stable flow was reached with the pump, the moving distance of the micro-beads were measured. The average flow velocity was half the micro-bead velocity at the centreline. The volumetric flow rates were calculated by multiplying the flow velocity with the cross-sectional area of the microfluidic channel. Each dot represents the mean ± standard deviation (n = 18).

**Supplementary Fig. S3.**
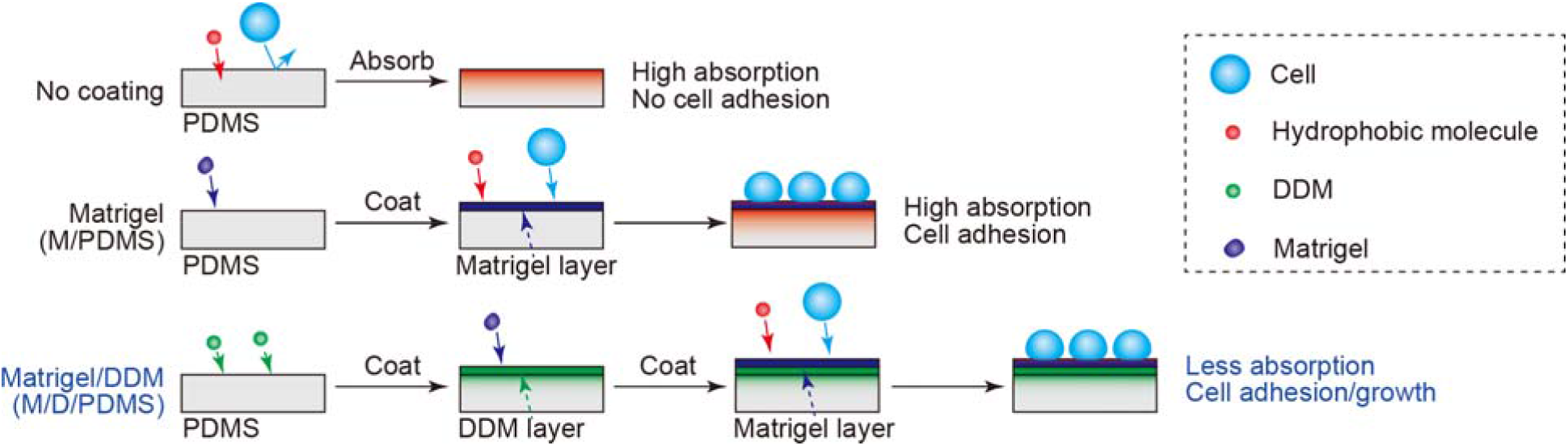
Schematic for PDMS coating with DDM and Matrigel to respectively prevent absorption of hydrophobic molecules in PDMS and promote cell adhesion and growth on PDMS.

**Supplementary Fig. S4.**
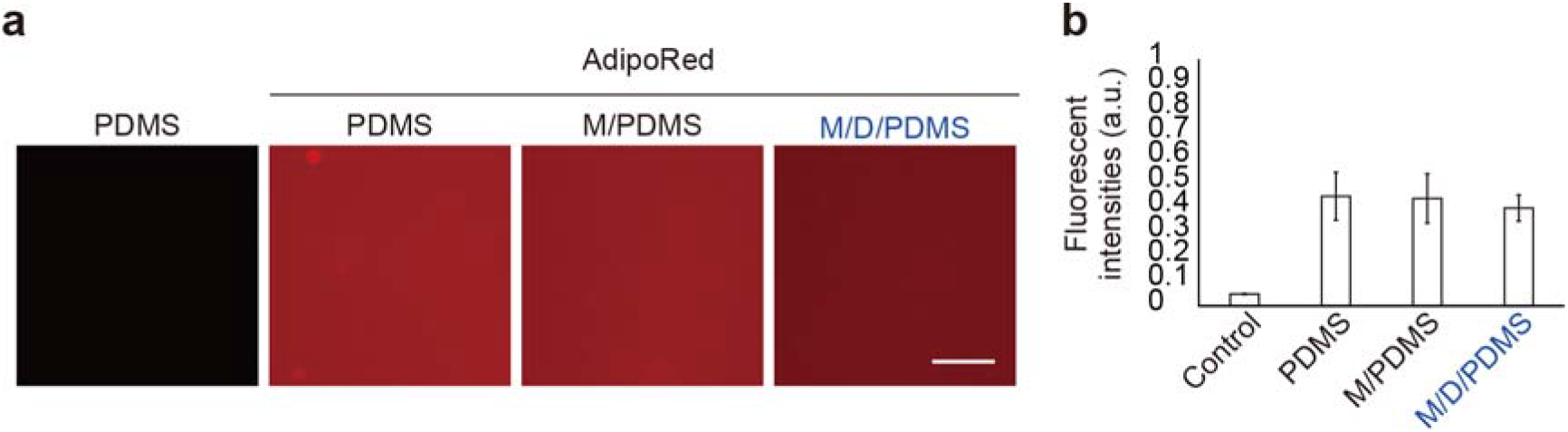
Absorption of hydrophobic molecules in PDMS. **a,** Fluorescent micrographs of PDMS coated with Matrigel (M/PDMS), Matrigel and DDM (M/D/PDMS), and PDMS without any coating (PDMS) that were treated with the AdipoRed lipid fluorescent dye. PDMS without any coating or treatment was used as the control. The scale bar represents 100 μm. **b,** Fluorescent intensities of AdipoRed dye absorbed in PDMS based on the microscopic photographs shown in **a**. The bars and error bars represent the mean ± standard deviations (n = 3).

**Supplementary Fig. S5.**
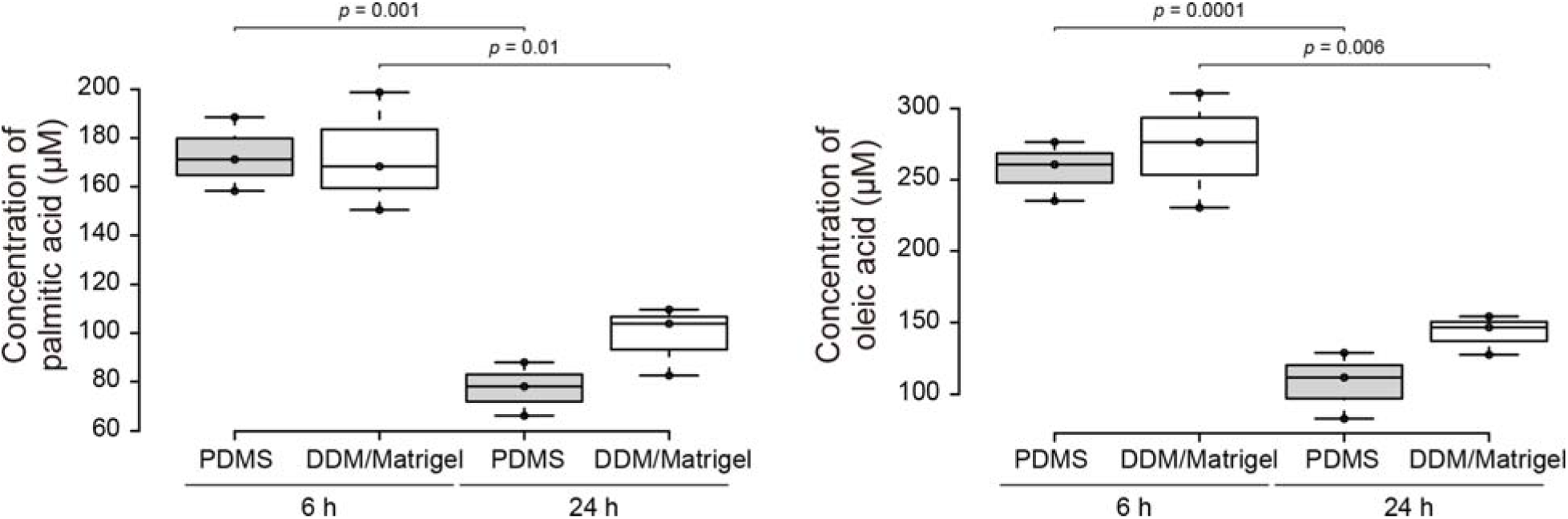
Lipid absorption in PDMS. FFA concentrations [left: palmitic acid (initial: 167 μM); right: oleic acid (initial: 333 μM)] in cell culture media after incubation in PDMS or DDM/Matrigel-coated microfluidic channels for 6 and 24 h. The centrelines show the medians. The box limits indicate the 25th and 75th percentiles. The whiskers extend 1.5 times the interquartile range from the maximum and minimum points; data points are plotted as circles. *p* values were estimated with Student’s *t*-test (n = 3).

**Supplementary Fig. S6.**
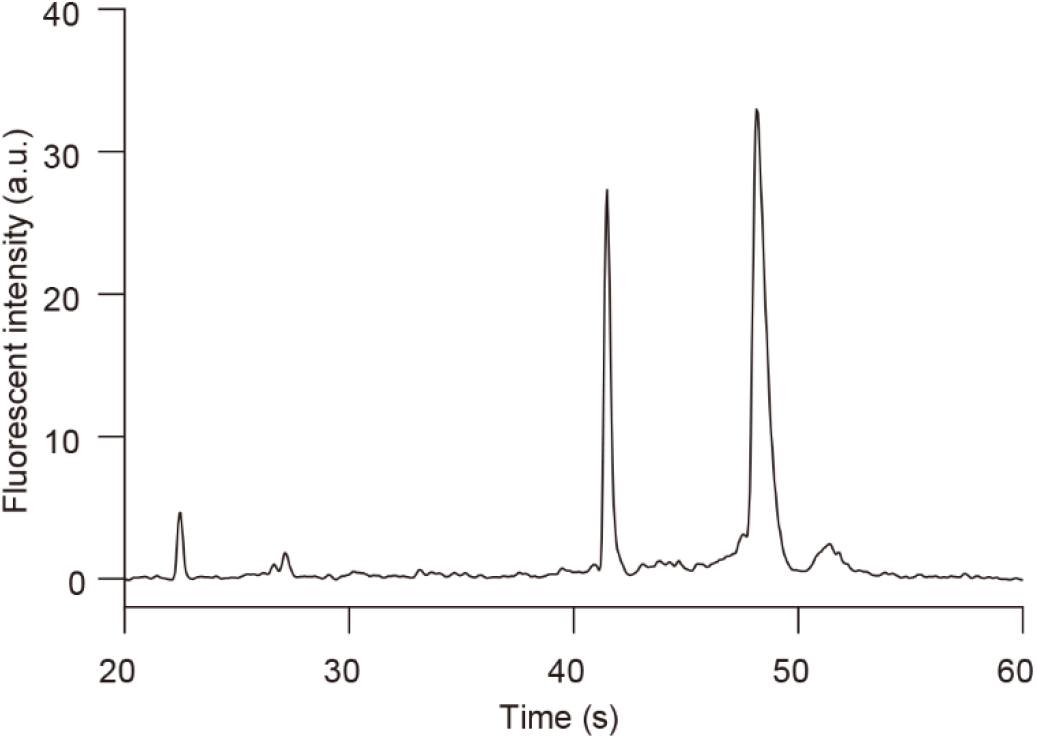
Typical electropherogram of total RNA obtained from Caco-2 cells cultured on a chip with Agilent 2100 Bioanalyzer. The electropherogram shows that 18S and 28S ribosomal RNA (rRNA) bands were clearly detected without smeared bands, and the RNA integrity number (RIN), which is used to standardise RNA quality control, was over 7.0. This result indicates that the total RNA harvested from the chip was sufficient to be applied to RNA sequencing.

**Supplementary Fig. S7.**
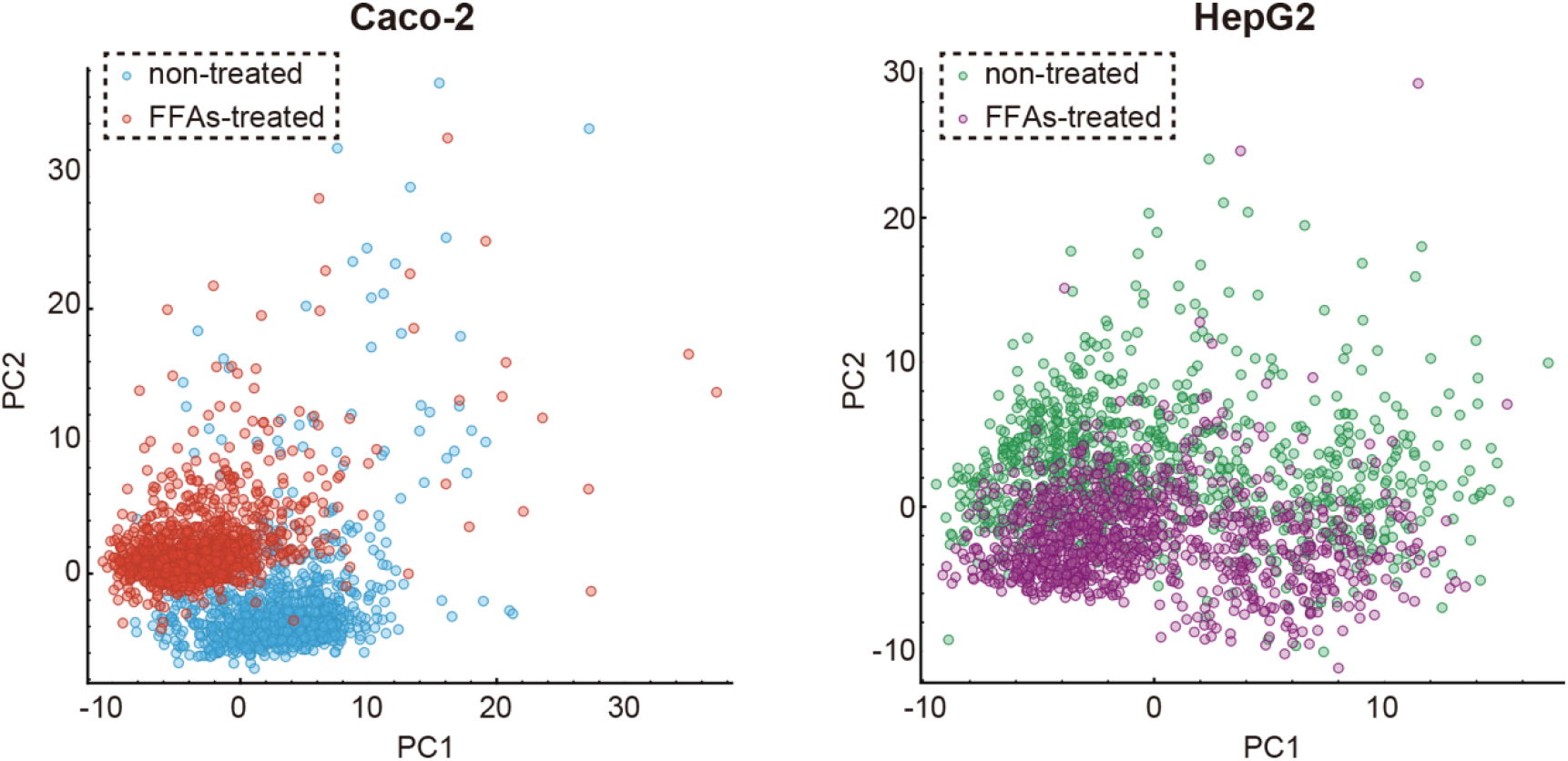
PCA for single-cell profiling of FFA-treated Caco-2 (*left*) and HepG2 (*right*) cells stained with Calcein-AM and Hoechst 33258. Compared with the t-SNE results shown in **Fig. 4c** and **4d,** PCA could not distribute cells in a two-dimensional graph, and most of the cells aggregated in a smaller area.

**Supplementary Fig. S8.**
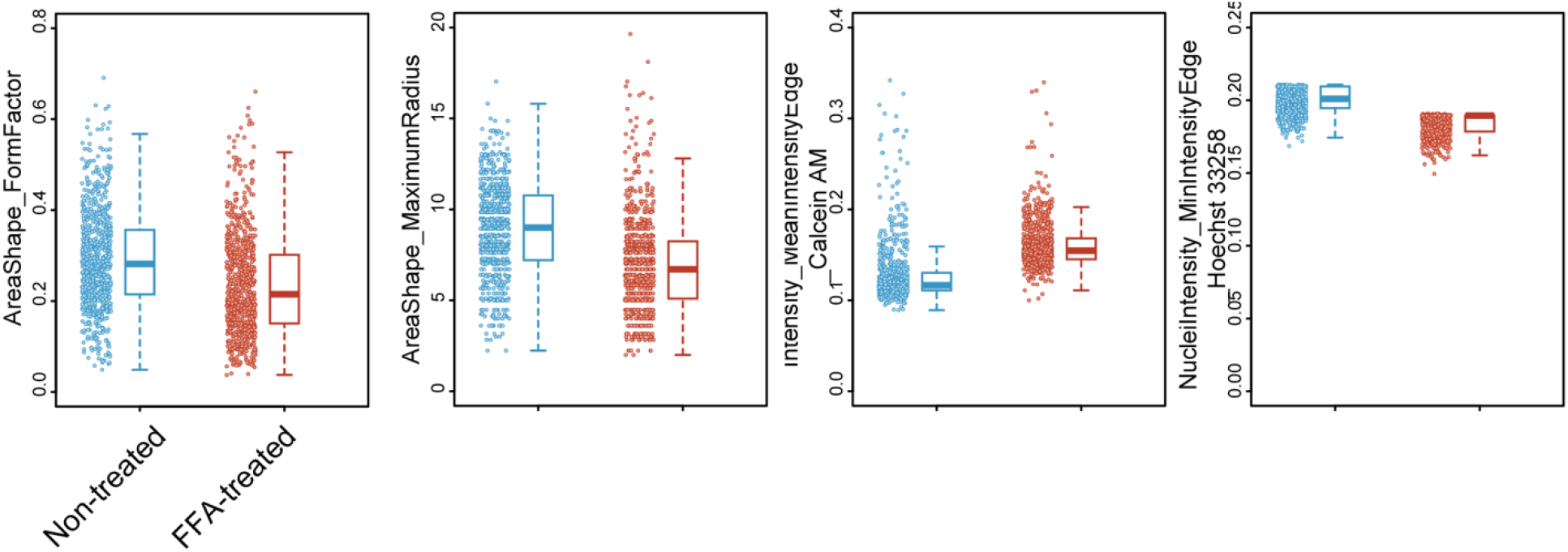
Boxplots comparing cellular parameters of non-treated and FFA-treated Caco-2 cells. The centrelines of the boxplots show the medians. The box limits indicate the 25th and 75th percentiles. The whiskers extend 1.5 times the interquartile range from the 25th and 75th percentiles. *p*-values were estimated with the asymptotic Wilcoxon signed-rank test and are presented in **Supplementary Table S8**.

**Supplementary Fig. S9.**
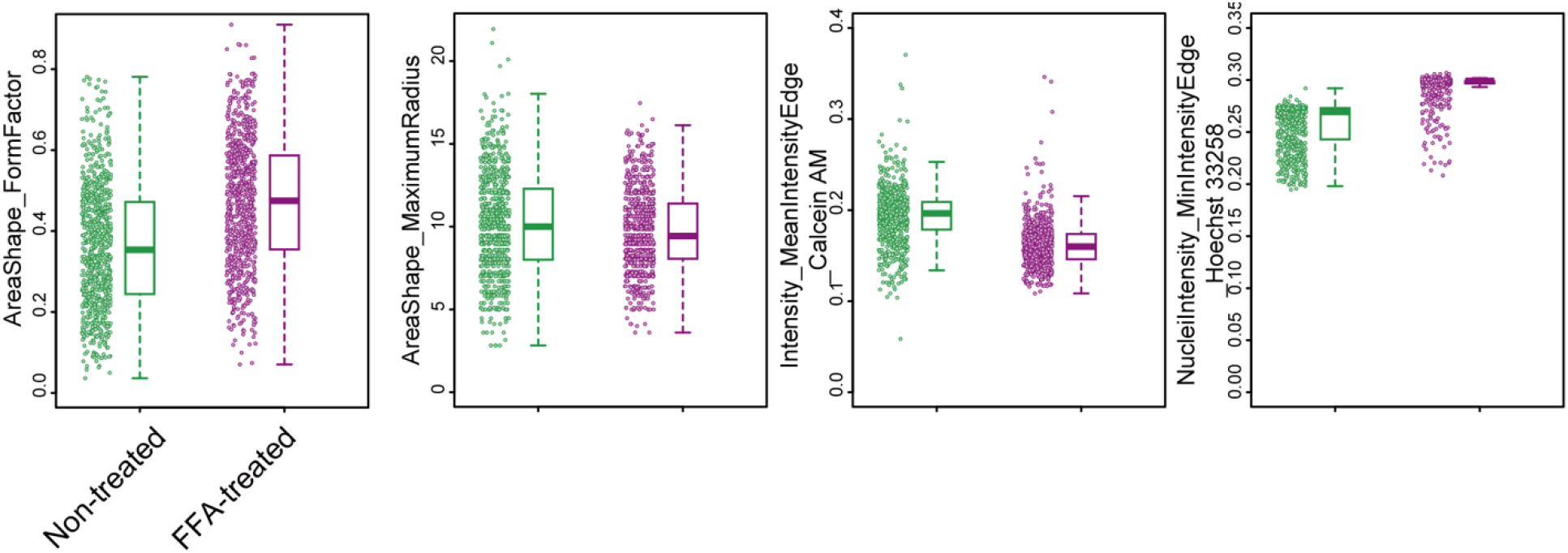
Boxplots comparing cellular parameters of non-treated and FFA-treated HepG2 cells. The centrelines of the boxplots show the medians. The box limits indicate the 25th and 75th percentiles. The whiskers extend 1.5 times the interquartile range from the 25th and 75th percentiles. *p*-values were estimated with the asymptotic Wilcoxon signed-rank test and are presented in Supplementary Table S9.

